# Structure of heme *d*_1_-free *cd*_1_ nitrite reductase NirS

**DOI:** 10.1101/2020.02.12.945543

**Authors:** Thomas Klünemann, Wulf Blankenfeldt

**Affiliations:** Structure and Function of Proteins, Helmholtz Centre for Infection Research, Inhoffenstraße 7, Braunschweig, Niedersachsen, 38124, Germany; Institute for Biochemistry, Biotechnology and Bioinformatics, Technische Universität Braunschweig, Spielmannstraße 7, Braunschweig, Niedersachsen, 38106, Germany

## Abstract

A key step in anaerobic nitrate respiration is the reduction of nitrite to nitric oxide, which is catalysed by *cd*_1_ nitrite reductase NirS in e.g. the gram-negative opportunistic pathogen *Pseudomonas aeruginosa*. Each subunit of this homodimeric enzyme consists of a cytochrome *c* domain and an eight-bladed β-propeller that binds the uncommon isobacteriochlorin heme *d*_1_ as an essential part of its active site. Although NirS is mechanistically and structurally well studied, the focus of previous studies has been on the active, heme *d*_1_-bound form. The heme *d*_1_-free form of NirS reported here, representing a premature state of the reductase, adopts an open conformation with the cytochrome *c* domains moved away from each other with respect to the active enzyme. Further, movement of a loop around W498 seems to be related to a widening of the propeller, allowing easier access to the heme *d*_1_ binding side. Finally, a possible link between the open conformation of NirS and flagella formation in *P. aeruginosa* is discussed.

**Synopsis:** The crystal structure of heme *d*_1_-free *cd*_1_ nitrite reductase NirS from *Pseudomonas aeruginosa* has been determined and provides insight into a premature form of the enzyme.

## 1. Introduction

Denitrification is the stepwise reduction of nitrogen oxides to dinitrogen and is utilized by many bacteria to replace the terminal oxygen-dependent steps of the respiratory chain under anaerobic conditions. A highly regulated step is the reduction of nitrite to nitric oxide, since both of these molecules are toxic to the cell. Many bacteria utilise the *cd*_1_ nitrite reductase NirS to catalyse this step, and the NirS enzymes from *Pseudomonas aeruginosa* (*Pa*) and *Paracoccus pantotrophus* (*Pp*) are functionally and structurally well studied. Both chains of this homodimeric enzyme possesses a c-type cytochrome domain with a covalently attached heme *c* functioning as electron entry point and a C-terminal eight-bladed β-propeller that utilizes the uncommon bacteriochlorin heme *d*_1_ as an active site cofactor (Figure 1a) (Nurizzo *et al.*, 1997; Williams *et al.*, 1997). Both domains interact via an N-terminal arm, which swaps domains in the case of *Pa*-NirS (Williams *et al.*, 1997; Nurizzo *et al.*, 1997). Compared to other tetrapyrroles such as heme *b*, high affinity for anionic molecules such as nitrite and low affinity for nitric oxide has been observed for the ferrous isobacteriochlorine heme *d*_1_. This is rooted in the unique carbonyl moieties at ring A and B and a double bond in the propionate sidechain at ring C (Chang *et al.*, 1986; Rinaldo *et al.*, 2011; Fujii *et al.*, 2016). During its reduction, nitrite is coordinated by H327 and H369 on top of the ferrous iron of heme *d*_1_ (Figure 1b)(Cutruzzola *et al.*, 2001). After dehydration and reduction of the substrate, the tetrapyrrole heme *d*_1_ is then reduced via internal electron transfer to allow efficient replacement of the product nitric oxide by another nitrite anion (Rinaldo *et al.*, 2011). In case of *Pa*-NirS, the intramolecular electron transfer is the rate limiting step and is allosterically controlled by smaller conformational changes within the cytochrome *c* domain (Farver *et al.*, 2009; Nurizzo *et al.*, 1999, 1998). Interestingly, when *Pp*-NirS was crystallised under reducing conditions, a ∼60° rotation of the cytochrome *c* domain around the pseudo eight-fold axis of the β-propeller was observed (Sjögren & Hajdu, 2001). This rotation was also seen in *Pa*-NirS after mutation of H327 to alanine (Brown *et al.*, 2001).

**Figure 1.**
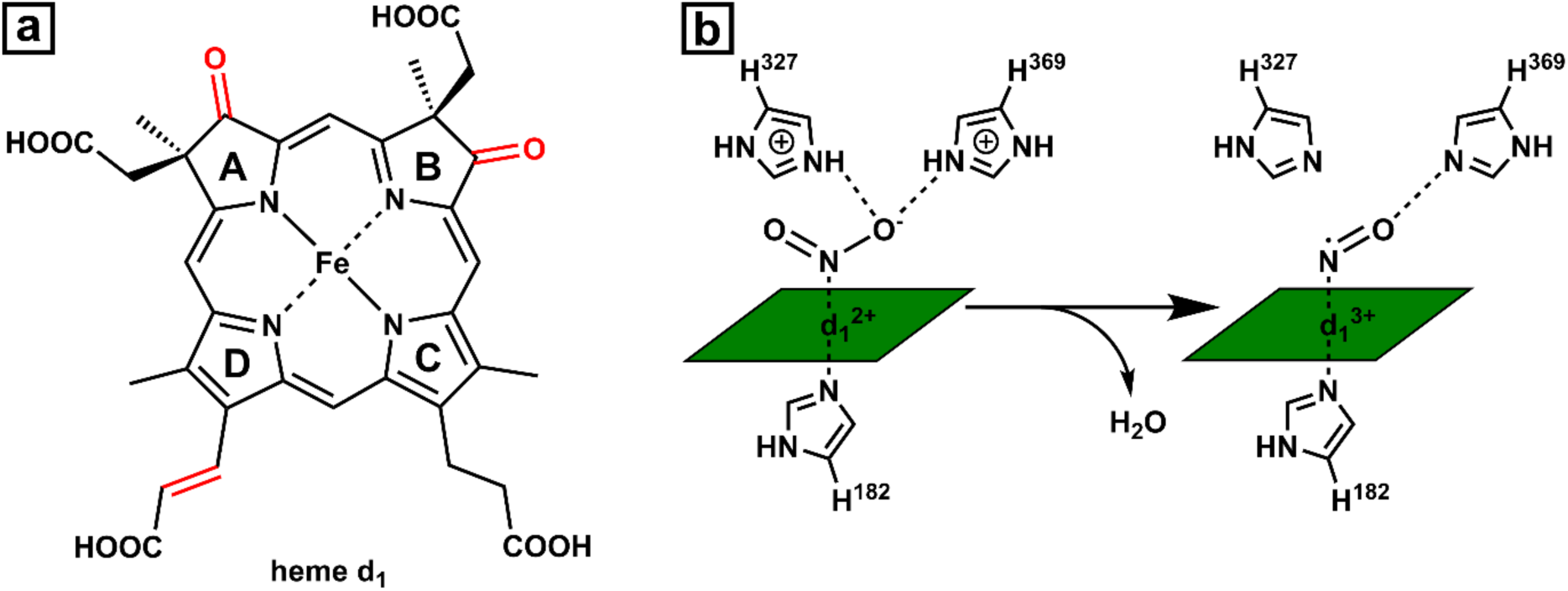
(a) Chemical structure of heme *d*_1_ with characteristic features highlighted in red. (b) Simplified depiction of the *Pa*-NirS active site during the reduction of nitrite to nitric oxide.

All structural studies of NirS up to now have focused on the heme *d*_1_-bound form of the enzyme and have led to a detailed understanding of the reaction mechanism within the denitrification context. Recently however, following the discovery that *Pa*-NirS forms complexes with the flagella protein FliC and the chaperone DnaK, an enzyme activity-independent scaffolding function of a heme *d*_1_-free form of NirS in flagella formation has been proposed (Borrero-de Acuña *et al.*, 2015). This cofactor-less form is also expected to exist in the course of maturation of NirS, where the enzyme has been demonstrated to transiently interact with NirF and NirN, two proteins involved in the biosynthesis of the heme *d*_1_ cofactor (Nicke *et al.*, 2013). We have therefore set out to determine the heme *d*_1_-free structure of NirS in order to gain more insight into these processes.

## 2. Materials and methods

### 2.1. Macromolecule production

NirS with bound dihydro-heme *d*_1_ was isolated from *Pseudomonas aeruginosa* strain RM361 (nirN::tet) (Kawasaki *et al.*, 1997) after cultivation under anaerobic conditions as previously described (Adamczack *et al.*, 2014; Klünemann *et al.*, 2019). Briefly, purification of NirS with bound dihydro-heme *d*_1_, the precursor of heme *d*_1_ that accumulates in the absence of NirN, was achieved by two subsequent ion exchange chromatographic steps as first described by Parr and colleagues (Parr *et al.*, 1976). The purified protein was dialysed against 10 mM Tris/HCl pH 8 and loaded onto a Q-Sepharose fast flow anion exchange column. After washing for 4 h with dialysis buffer at a flowrate of 1 ml/min the eluted protein lost dihydro-heme *d*_1_ but still contained the covalently attached heme *c* as indicated by UV/Vis absorption band at 410 nm (indicative of heme *c*) and no absorption at 630 nm (indicative of dihydro-heme *d*_1_) during elution. Interestingly, dihydro-heme *d*_1_ does not seem to be eluted during the washing procedure but remained bound to the resin, visible as a greenish coloration of the chromatography column. Prior to crystallisation, the protein was subjected to size exclusion chromatography (Superdex 200, GE Healthcare) in a buffer containing 10 mM Tris/HCl pH 8 with 150 mM NaCl.

### 2.2. Crystallization

Crystallisation was performed by vapour diffusion in sitting drop 96-3 Intelli plates (Art Robbins Instruments) at room temperature. Plates were set up with a with a HoneyBee 961 pipetting robot (Digilab Genomic Solutions), which mixed 200 nl of protein solution with 200 nl of mother liquor and provided a precipitant reservoir of 60 µl. To get structural insight into possible changes in the dihydro-heme *d*_1_ bound form, a screen based on previously published crystallisation conditions for NirS (Tegoni *et al.*, 1994) (Brown *et al.*, 2001) was performed. After two days, plate-shaped green crystals were observed with phosphate as the precipitant (1.9 M K_2_HPO_4_ and 0.1 M Tris-HCl pH 8). The green colour hinted at the presence of dihydro-heme *d*_1_ in these crystals (Figure 2). Surprisingly, under conditions with PEG4000 or PEG6000, red rod-shaped or tetragonal crystals started to grow. The red coloration indicates that dihydro-heme *d*_1_ was lost as a cofactor. As these crystals only diffracted x-ray to 4 Å, sparse matrix screening with the commercially available crystallization suites JCSG+ (Quiagen) and Morpheus (Molecular Dimensions) was used to identify conditions better suited for diffraction experiments. The best-diffracting crystals for heme *d*_1_-free NirS occurred with 0.1 M phosphate/citrate pH 4.2 and 40% (v/v) PEG300, with dihydro-heme *d*_1_ removed from NirS by the chromatographic procedure described above prior to crystallization experiment. Crystals were flash frozen in liquid nitrogen after cryo-protection with 10% (v/v) (*R,R*)-2,3-butandiol.

**Figure 2.**
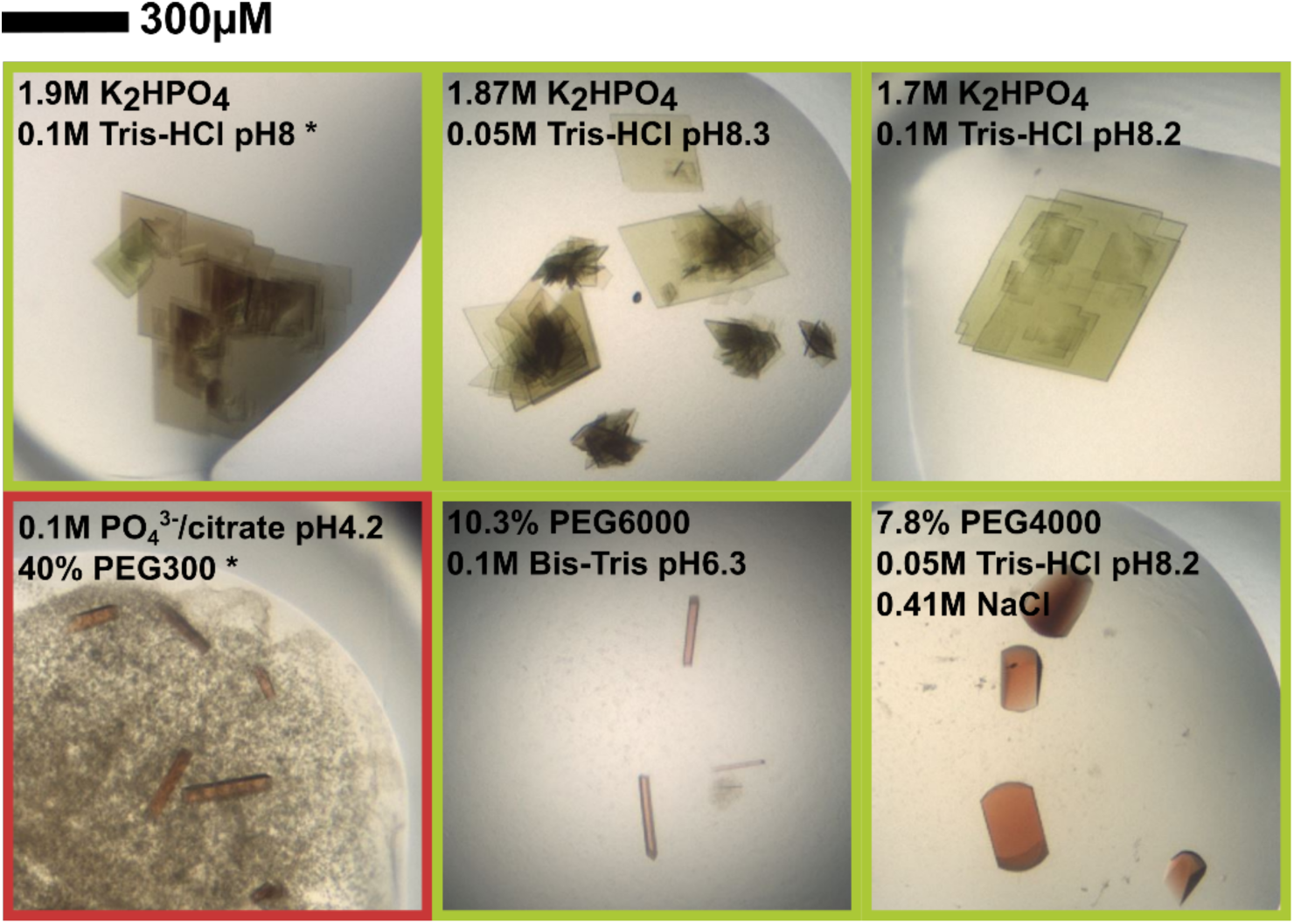
Photographs of representative crystals grown from NirS isolated with (green frame) or without (red frame) bound dihydro-heme *d*_1_. The data sets used in this study were collected from crystallization conditions marked with an asterisk (*)

### 2.3. Data collection and processing

3600 diffraction images, each with an oscillation angle of 0.1°, were collected at beamline P11 at PETRA III (DESY, Hamburg, Germany) on a PILATUS 6M fast detector. Images were processed utilizing the autoPROC pipeline (Vonrhein *et al.*, 2011) executing XDS (Kabsch, 2010), Pointless (Evans, 2011) and Aimless (Evans & Murshudov, 2013). In case of NirS with bound dihydro-heme *d*_1_ the crystals diffracted anisotropically, which was accounted for by using STARANISO (Tickle *et al.*, 2018) within autoPROC.

**Table 1.** Data collection and processing. Values for the outer shell are given in parentheses.

|  | heme *d*<sub>1</sub> free NirS | dihydro-heme *d*<sub>1</sub> bound NirS |
| --- | --- | --- |
| Diffraction source | P11 at PETRAIII | P11 at PETRAIII |
| Wavelength (Å) | 1.033 | 1.739 |
| Temperature (K) | 100 | 100 |
| Detector | Pilatus 6M fast | Pilatus 6M fast |
| Crystal-detector distance (mm) | 368.1 | 168.4 |
| Rotation range per image (°) | 0.1 | 0.1 |
| Total rotation range (°) | 360 | 360 |
| Exposure time per image (s) | 0.1 | 0.05 |
| Space group | $P4_322$ | $P2_12_12$ |
| $a, b, c$ (Å) | 67.25, 67.25, 277.10 | 163.76, 91.09, 112.71 |
| $\alpha, \beta, \gamma$ (°) | 90, 90, 90 | 90, 90, 90 |
| Resolution range (Å) | 48.25–1.86 (1.93–1.86) | 112.71–2.38 (2.62–2.38) |
| Resolution limits along axes ($a^*$, $b^*$, $c^*$) | - | 2.89, 2.38, 2.51 |
| Total No. of reflections | 1363122 (93274) | 625785 (23996) |
| No. of unique reflections | 54709 (5373) | 50095 (2505) |
| Completeness spherical (%) | 99.97 (100) | 73.1 (14.7) |
| Completeness ellipsoidal (%) | - | 92.6 (44.5) |
| Redundancy | 24.9 (17.4) | 12.5 (9.6) |
| $\langle I/\sigma(I) \rangle$ | 20.3 (1.8) | 6.6 (1.6) |
| CC<sub>1/2</sub> | 0.99 (0.57) | 0.99 (0.64) |
| $R_{p.i.m.}$ | 0.03 (0.32) | 0.11 (0.54) |
| Overall $B$ factor from Wilson plot (Å<sup>2</sup>) | 26.2 | 23.6 |

### 2.4. Structure solution and refinement

The crystal structure of heme *d*_1_-free NirS was determined by molecular replacement utilizing Phaser (McCoy *et al.*, 2007) and coordinates of the H327A variant of *Pa*-NirS (NirS^H327A^; PDB:1hzu (Brown *et al.*, 2001)), which crystalized in the same space group but with approx. 3 Å longer unit cell axes. The structure of NirS with bound dihydro-heme *d*_1_ was phased by Fourier synthesis using phases from a published structure of *Pa*-NirS with bound heme *d*_1_ (PDB: 1nir (Nurizzo *et al.*, 1997)). To remove phase bias, both initial models were subjected to ten cycles of refinement with enabled simulated annealing in phenix.refine (Afonine *et al.*, 2012). The final models were constructed by several cycles of manual optimization in Coot (Emsley & Cowtan, 2004) and computational refinement including TLS refinement. Depictions of the final models were prepared with PyMol (Schrödinger, 2015).

**Table 2.** Structure solution and refinement. Values for the outer shell are given in parentheses.

| | heme $d_1$ free NirS | dihydro-heme $d_1$ bound NirS |
| --- | --- | --- |
| Resolution range (Å) | 48.25–1.86 (1.93–1.86) | 79.61–2.38 (2.42–2.38) |
| Final $R_{\text{cryst}}$ | 0.17 (0.23) | 0.24 (0.33) |
| Final $R_{\text{free}}$ | 0.19 (0.27) | 0.28 (0.43) |
| No. of non-H atoms |  |  |
| Protein | 4100 | 8364 |
| Ligand | 65 | 214 |
| Water | 255 | 246 |
| R.m.s. deviations |  |  |
| Bonds (Å) | 0.01 | 0.01 |
| Angles (°) | 0.94 | 1.37 |
| Average $B$ factors (Å<sup>2</sup>) | 31.0 | 23.6 |
| Ramachandran plot |  |  |
| Most favoured (%) | 96.35 | 94.24 |
| Allowed (%) | 3.65 | 5.67 |

## 3. Results

In previous work with dihydro-heme *d*_*1*_ dehydrogenase NirN, we isolated the *cd*_*1*_ nitrite reductase NirS with bound dihydro-heme *d*_1_, the final intermediate of heme *d*_1_ biosynthesis, which differs from heme *d*_1_ by missing a double bond in the propionate chain attached to ring D in comparison to heme *d*_1_ itself (Klünemann *et al.*, 2019). It has previously been shown that NirS can utilize both cofactors but is less active with dihydro-heme *d*_1_ (Hasegawa *et al.*, 2001; Kawasaki *et al.*, 1997; Adamczack *et al.*, 2014). To determine if the altered activity is caused by structural changes or by inherent properties of dihydro heme *d*_1_, NirS with bound dihydro heme *d*_*1*_ was crystalized with conditions published for heme *d*_1_-bound NirS (Tegoni *et al.*, 1994; Brown *et al.*, 2001). In this complex, anomalous difference density around the iron centres indicates similar occupancies for dihydro-heme *d*_1_ and heme c, the latter being covalently attached to the polypeptide. Comparison with the previously published heme *d*_1_-bound structure of NirS reveals no significant differences (Nurizzo *et al.*, 1997)(C_α_-rmsd: 0.38 Å; Figure 3+4). This suggests that the reported differences in the activity of NirS loaded with dihydro-heme *d*_1_ or heme *d*_1_ likely are caused by inherent properties of the cofactor.

**Figure 3.**
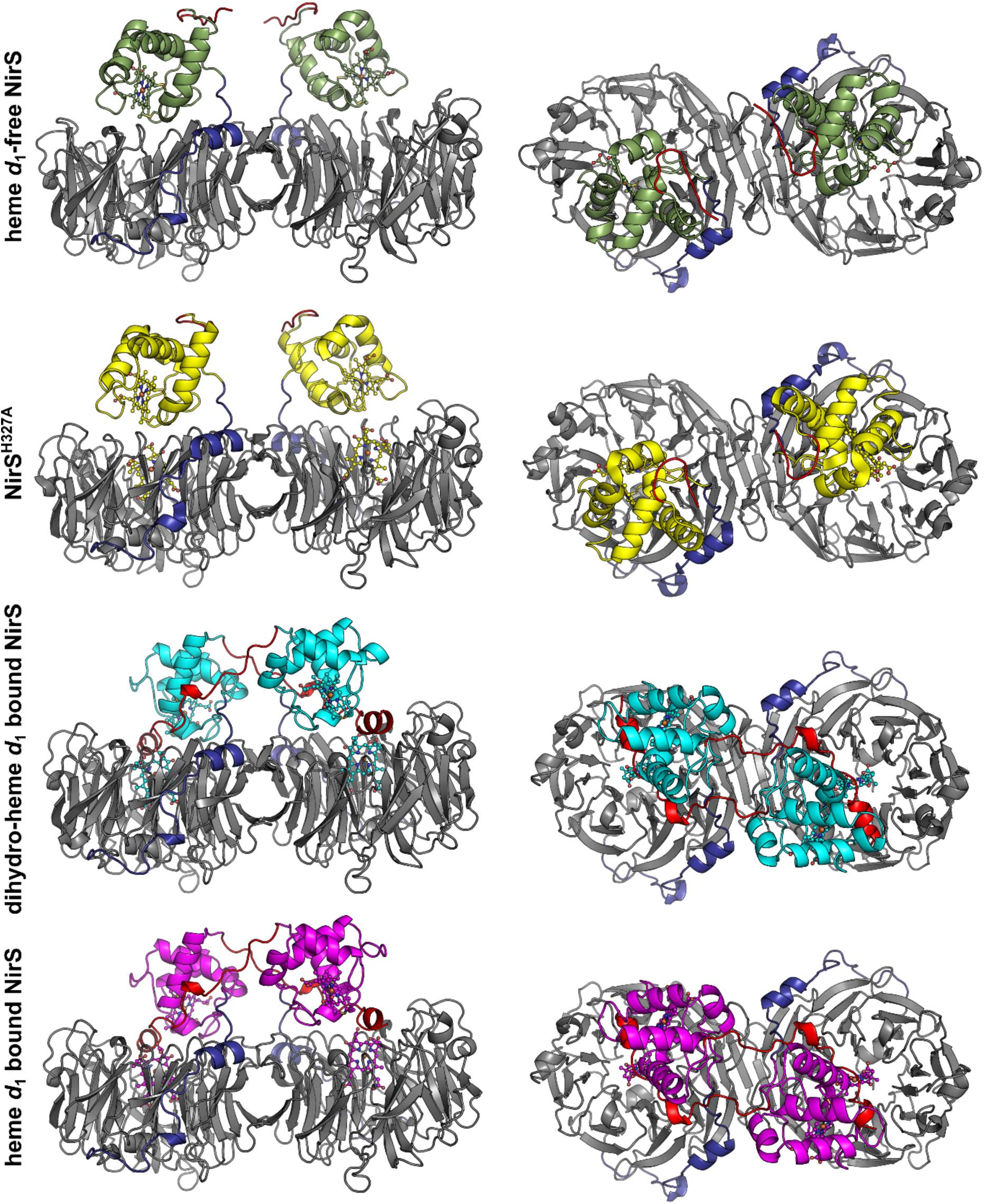
Cartoon representation of different conformation observed for *cd*_1_ nitrite reductase NirS of *P. aeruginosa* (heme *d*_1_ bound NirS: 1nir (Nurizzo *et al.*, 1997); NirS^H327A^: 1hzu (Brown *et al.*, 2001)) after superposition of the *d*_1_-domains. The N-terminal arm is coloured in red, the linker in blue and the *d*_1_-domain in grey. The cytochrome *c* domain and the tetrapyrroles are coloured individually.

**Figure 4.**
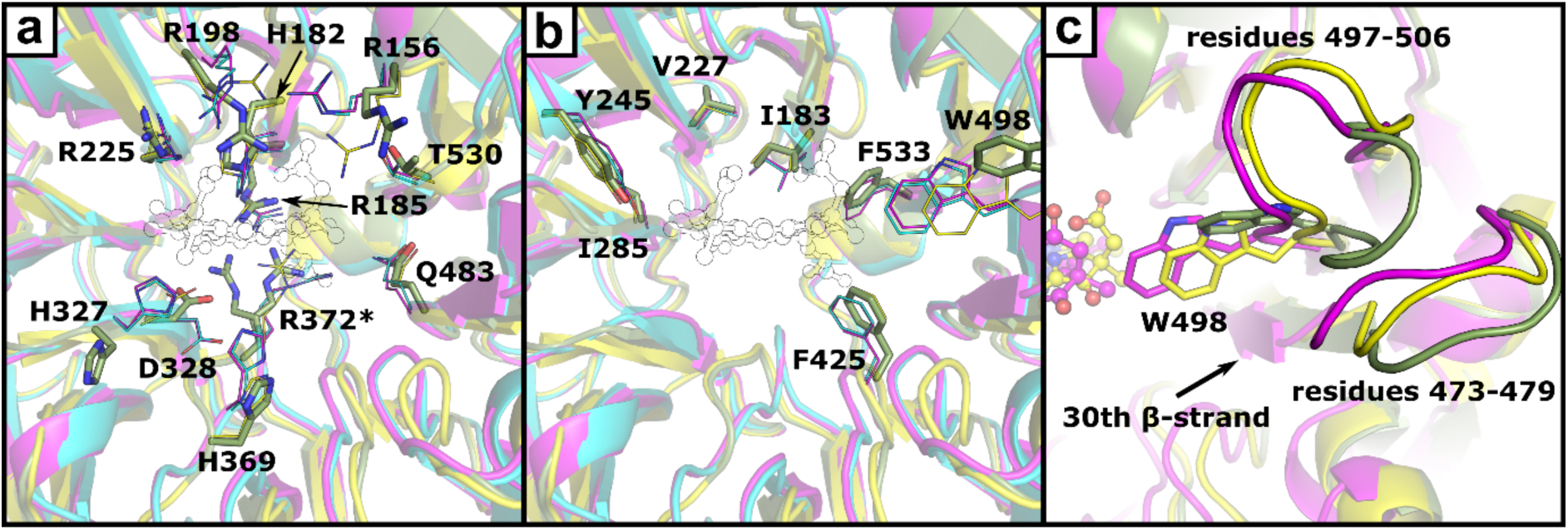
Depiction of hydrophilic (a) and hydrophobic (b) residues in the heme *d*_1_ binding pocket. Amino acids are shown as sticks (heme *d*_1_-free NirS, green) or lines (heme *d*_1_ bound NirS, magenta; NirS^H327A^, yellow; dihydro-heme *d*_1_ bound NirS, cyan). Residues with alternativ conformations are marked with an asterisk (*). The position of heme *d*_1_ inside the binding pocket is shown as a thin outline. Panel c depicts the movement of the loops associated with W498 narrowing the binding pocket of heme *d*_1_-free NirS after binding a *d*_1_-type hemes.

The structure of heme *d*_1_-free NirS revealed a similar dimeric structure and domain organization as mature, heme *d*_1_-bound NirS (Nurizzo *et al.*, 1997). Both chains have a C-terminal eight-bladed β-propeller, which has previously been termed the *d*_1_-domain as it is responsible for binding *d*_1_-type tetrapyrroles in NirS and the related proteins NirF and NirN (Figure 3). In NirS, these domains form homodimers by interaction of the 15^th^ β-strand of each subunit, which is consistent with the SEC-data collected during purification (data not shown). The N-terminal domain consists of a cytochrome *c* fold that is connected *via* a linker domain to the *d*_1_-domain and harbours a covalently attached heme *c*. Interestingly, the position of the cytochrome *c* domain differs from the one found in the heme *d*_1_ and dihydro-heme *d*_1_-bound structures by a rotation of approximately 60° around the pseudo eight-fold axis of the β-propeller. A similar conformation has already been observed for *Pa*-NirS after amino acid exchange of H327 to alanine or for *Pp*-NirS crystallised under reducing conditions (Brown *et al.*, 2001; Sjögren & Hajdu, 2001). No density was observed for the N-terminal arm, which resides between the cytochrome c domain and the *d*_1_-domain of the opposing subunit in NirS in the closed conformation, indicating that it is flexible in the heme *d*_1_-free form of the enzyme.

An overlay of the cytochrome *c* domain of heme *d*_1_-free NirS with other *Pa*-NirS structures reveals higher similarity with the structure of NirS reduced *in crystallo* (C_α_ rmsd: 0.37 Å) as compared to the oxidised form (C_α_ rmsd: 1.09 Å) (Nurizzo *et al.*, 1998; Nurizzo *et al.*, 1997). The difference is mostly caused by the position of residues 55-64 and consistent with observation previously discussed for NirS^H327A^ (C_α_ rmsd cytochrome *c*: 0.35 Å). A similar comparison of the *d*_1_-domain reveals a lower C_α_ rmsd with respect to the H327A variant of *Pa*-NirS (0.75 Å) than to the oxidised or reduced wildtype protein (1.34 Å and 1.39 Å). NirS^H327A^ has been shown to have a wider β-propeller, allowing better solvent access to heme *d*_1_ (Brown *et al.*, 2001). Further, the reported occupancy (0.5) of heme *d*_1_ together with the observed change of coloration of these crystals in the course of time, indicative for the loose of heme *d*_1_, suggests that the open conformation allows easy access to the ligand binding side in general (Brown *et al.*, 2001). Closer inspection of the heme *d*_1_ binding pocket reveals only few distinct changes related to *d*_1_-type heme binding (Figure 4). Three arginines that interact with the acetate or propionate sidechains of heme *d*_1_ in the heme *d*_1_ bound structure (R156, R198, R372) adopt different conformations in the heme *d*_1_-free form. Further, H327 and H369, which are both involved in substrate binding and catalysis by NirS, have different positons. Interestingly, H182, which acts as the fifth ligand to the iron cation in the centre of heme *d*_1_, shows no conformational changes without bound tetrapyrrole. This stands in contrast to other d_1_-type heme binding β-propellers of the heme *d*_1_ biosynthesis proteins NirN and NirF, where the corresponding histidine rotates with respect to the ligand-free form to coordinate the iron (*Klünemann et al*., under revision; DOI: 10.1101/2020.01.13.904656) (Klünemann *et al.*, 2019). Another distinctive variation in the *d*_1_-domain of heme *d*_1_-free and bound NirS is observed for the loop consisting of residues 497-506, which moves upon heme *d*_1_ binding to enable W498 to lock the cofactor inside its binding pocket. Interestingly, the neighbouring loop (residues 473-479) also takes a slightly different conformation. This loop is connected to the 30^th^ β-strand, which is inside of the propeller and moved inward upon heme *d*_1_ binding, indicating a connection between the propeller widening and the movement of W498 (Fig. 3c).

## 4. Discussion

It has long been known that the *cd*_1_ nitrite reductase NirS adopts an open and a closed conformation, and it has been hypothesized that these states are linked to the catalytic cycle (Brown *et al.*, 2001; Sjögren & Hajdu, 2001). The data presented here, however, suggest that the open conformation rather is related to a native but immature form of the enzyme and is therefore not necessarily associated to the catalytic cycle. This is further supported by the observation that *Pp*-NirS is unable to undergo a full catalytic cycle *in crystallo* when crystallised in the open conformation (Sjögren & Hajdu, 2001). In addition, a complex between *Pa-*NirS in its closed conformation and the nitric oxide reductase (NOR) has been published recently, suggesting an association between NirS and the cytoplasmic membrane that enables substrate channelling (Terasaka *et al.*, 2017). The open conformation would lead to a change of the membrane interface of NirS and therefore cannot support this membrane association, indicating that it is not part of the catalytic cycle.

The open conformation of NirS may, however, have additional physiological importance outside of the denitrification process. It has recently been found that NirS can function as scaffold protein for flagella formation under denitrifying conditions, even in the absence of heme *d*_1_ (Borrero-de Acuña *et al.*, 2015). Interestingly, the N-terminal arm and the cytochrome *c* domain of NirS were shown to be involved in complex formation between NirS, FliC and DnaK in this process (Borrero-de Acuña *et al.*, 2015). Further, kinetic studies with NirS from *Pseudomonas stutzeri*, which lacks the N-terminal arm, and mutation of a conserved tyrosine in *Pa*-NirS and *Pp*-NirS suggest that the N-terminal arm is not required for the enzymatic activity, despite its interaction with heme *d*_1_ (Cutruzzolà *et al.*, 1997; Gordon *et al.*, 2003; Wilson *et al.*, 2001). Therefore, the conformational change observed in our study may be envisioned to release the N-terminal arm to allow flagella formation, thereby regulating cell motility. Further investigation will be required to confirm that the conformational changes observed *in crystallo* can also be found in solution and is associated with distinct functions of NirS *in vivo*.

## Acknowledgements

We thank the beamline staff at P11 at the PETRAIII synchrotron (Deutsches Elektronensynchrotron DESY, Hamburg, Germany) for letting us use their facilities. This work was supported by grants from the Deutsche Forschungsgemeinschaft (PROCOMPAS graduate school, GRK 2223/1).

